# Temporal evolution of beta bursts in the parkinsonian cortico-basal ganglia network

**DOI:** 10.1101/458414

**Authors:** Hayriye Cagnan, Mallet Nicolas, Christian K.E. Moll, Alessandro Gulberti, Manfred Westphal, Christian Gerloff, Andreas K. Engel, Wolfgang Hamel, Peter J. Magill, Peter Brown, Andrew Sharott

## Abstract

Prevalence and temporal dynamics of transient oscillations in the beta frequency band (15-35 Hz), referred to as beta bursts, are correlated with motor performance and tactile perception. Disturbance of these activities is a candidate mechanism for motor impairment in Parkinson’s disease (PD), where the excessively long bursts correlate with symptom severity and are reduced by pharmacological and surgical treatments. To date, characterization of beta bursts in PD has been limited to the local field potentials in the subthalamic nucleus (STN) and cortical EEG. Here, we describe the changes that take place in spiking activity across the cortico-basal ganglia circuit, providing a unique insight into the network dynamics of these transient oscillations. Firstly, we demonstrate that rhythmic subthalamic spiking activity emerges at a fixed phase relationship with respect to cortical beta bursts in PD patients. Using multichannel recordings of ensembles of neurons in the 6-OHDA rat model of PD, we then dissect the beta burst dynamics across the sensorimotor cortex and several basal ganglia structures: striatum (Str), globus pallidus externus (GPe) and STN. Each subcortical structure exhibits enhanced rhythmic activity in the beta band locked to the onset of cortical beta bursts and longer cortical bursts lead to stronger subcortical rhythmicity. Crucially, enhanced subcortical rhythmic activity emerges at a fixed phase relationship with respect to the motor cortex, comparable to the relationship observed in PD patients. Striatal beta bursts terminate prior to the recruitment of those in the STN and GPe, suggesting that while they could play an important role in establishing synchrony in the beta band, they do not extensively contribute to its maintenance in other basal ganglia structures. Critically, changes in cortico-subcortical phase coupling precede the onset of a cortical beta burst, supporting the hypothesis that phase alignment across the cortico-basal ganglia network could recruit these structures into synchronous network oscillations. We provide a powerful approach that not only examines pathophysiology of PD across the motor circuit, but also offer insights that could aid in the design of novel neuromodulation strategies to manipulate the state of the motor system before pathological activities emerge.

## Introduction

Oscillations in the beta frequency band (15-35 Hz) have been the focus of numerous studies due to their central role in cortical processing, particularly in mediating top-down commands (Bastos et al., 2015; Engel and Fries, 2010; Michalareas et al., 2016; Richter et al., 2017), for their prominence in the motor system (Baker, 2007; Baker et al., 1997; Klostermann et al., 2007; Sanes and Donoghue, 1993) and during Parkinson’s disease (PD) (Cassidy et al., 2002; Kühn et al., 2004; Lalo et al., 2008). Across different experimental conditions, changes in beta band activity have been linked with motor performance (Kühn et al., 2004; Tan et al., 2014), decision-making (Herz et al., 2018), motor-task completion (Feingold et al., 2015), tactile perception (Shin et al., 2017) and PD treatment state (Kühn et al., 2006, 2009; Little et al., 2013; Tinkhauser et al., 2017a, 2017b).

Several theories have been put forward regarding the origin of beta oscillations in the corticobasal ganglia (BG) circuit such as (1) the excitatory-inhibitory coupling between the subthalamic nucleus (STN) and the external segment of the globus pallidus (GPe) (Holgado et al., 2010; Terman et al., 2002), (2) cholinergic interneurons in the striatum (Str) (McCarthy et al., 2011) (3) cortical laminar dynamics (Roopun et al., 2006), and (4) subcortical input modulating cortical activity in a laminar specific fashion (Sherman et al., 2016). However, beta oscillations in the entire cortico-BG circuit are suppressed in PD when the STN is lesioned, or when continuous high frequency deep brain stimulation (DBS) is applied to either STN or the internal segment of the globus pallidus, inducing a reversible information-lesion (Eusebio et al., 2011; Kühn et al., 2008; Rosin et al., 2011). These observations highlight that presence of neural activity in the beta frequency band strongly depends on the distributed network architecture of the motor circuit in PD.

Until recently, changes in rhythmic activity patterns have been reported as an average state change, modulated by either the experimental condition or patients’ treatment state. However, recent studies have revealed that beta band activity occurs in transient bursts (Feingold et al., 2015; Sherman et al., 2016; Tinkhauser et al., 2017a, 2017b). While adaptive DBS (i.e. high frequency stimulation delivered only when beta power exceeds a certain threshold), and dopaminergic medication shift the distribution of beta burst durations towards shorter durations in the STN, continuous DBS suppresses all transient beta oscillations regardless of burst duration (Tinkhauser et al., 2017b). The relationship between treatment state and beta burst duration raises the possibility that it is not the presence of beta oscillations that is pathophysiological in PD, but the sustained propagation of the burst in the cortical-BG circuit (Cagnan et al., 2015; Tinkhauser et al., 2017a, 2017b, 2018). However, it remains unknown what the downstream effects of beta bursts are on spiking activity in different structures and how their duration modulates communication between different neuronal populations. Accurate characterization of neural dynamics surrounding a beta burst has the potential to guide the next generation of closed-loop DBS strategies, with the aim of preventing the generation and/or propagation of pathological network activities.

In this study, we (1) demonstrate that cortical beta bursts are coupled to transient rhythmicity in STN spiking activity in PD patients, (2) characterize how transient beta oscillations are relayed across different BG nuclei to accurately determine neural dynamics associated with the PD pathophysiology, (3) determine how cortico-BG interactions could contribute to the time-course of neural entrainment in different BG nuclei.

## Materials and Methods

### Intraoperative recordings

#### Patient details and clinical scores

The present study was conducted in agreement with the Code of Ethics of the World Medical Association (declaration of Helsinki, 1967), and received local ethics approval. Written informed consent was given by all patients who participated in this study. Cortical EEGs and microelectrode recordings from the STN were recorded from 7 patients (4 female, 3 male; age: 67 ± 3 years) with advanced idiopathic PD with a disease duration of 17 ± 8 years, during bilateral implantation of DBS electrodes in the STN, guided by microelectrode mapping. Clinical details are summarised in Supplemental Table 1.

#### Microelectrode recordings

Microelectrode recordings were acquired from five electrodes (Figure 1A) (MicroGuide, Alpha-Omega, Nazareth, Israel). While the central electrode aimed at the theoretical target (Supplementary Materials), the remaining four electrodes (FHC Inc., Bowdoinham, ME, USA) were placed 2mm around the central one. Signals were amplified (x20.000), sampled at 24 kHz, and filtered between 300 to 6000 Hz. Unit activities were classified as subthalamic unit activity using several previously described criteria: a well-defined elevation in the background activity (Moran et al., 2006; Sharott et al., 2014) together with presence of discharge patterns with tonic irregular, oscillatory or burst-firing features. These features allowed for a distinction to be made from the neighbouring substantia nigra pars reticulata thalamus/zona incerta neurons.

#### Cortical EEG signals

Cortical EEG signals were recorded, amplified and filtered (x4.000; bandpass, 0-400 Hz) using the 32-channel AlphaLab system (Alpha Omega Inc., Nazareth, Israel). Signals were recorded at the same time as the microelectrode recordings (Moll et al., 2015) and were sampled at 3005 Hz. EEGs were recorded approximately from the Fz and Cz using Ag/AgCl cup electrodes filled with conductive gel (Nicolet Biomedical, Madison, WI, USA). In this study, the Fz was referenced to the Cz.

**Figure 1.**
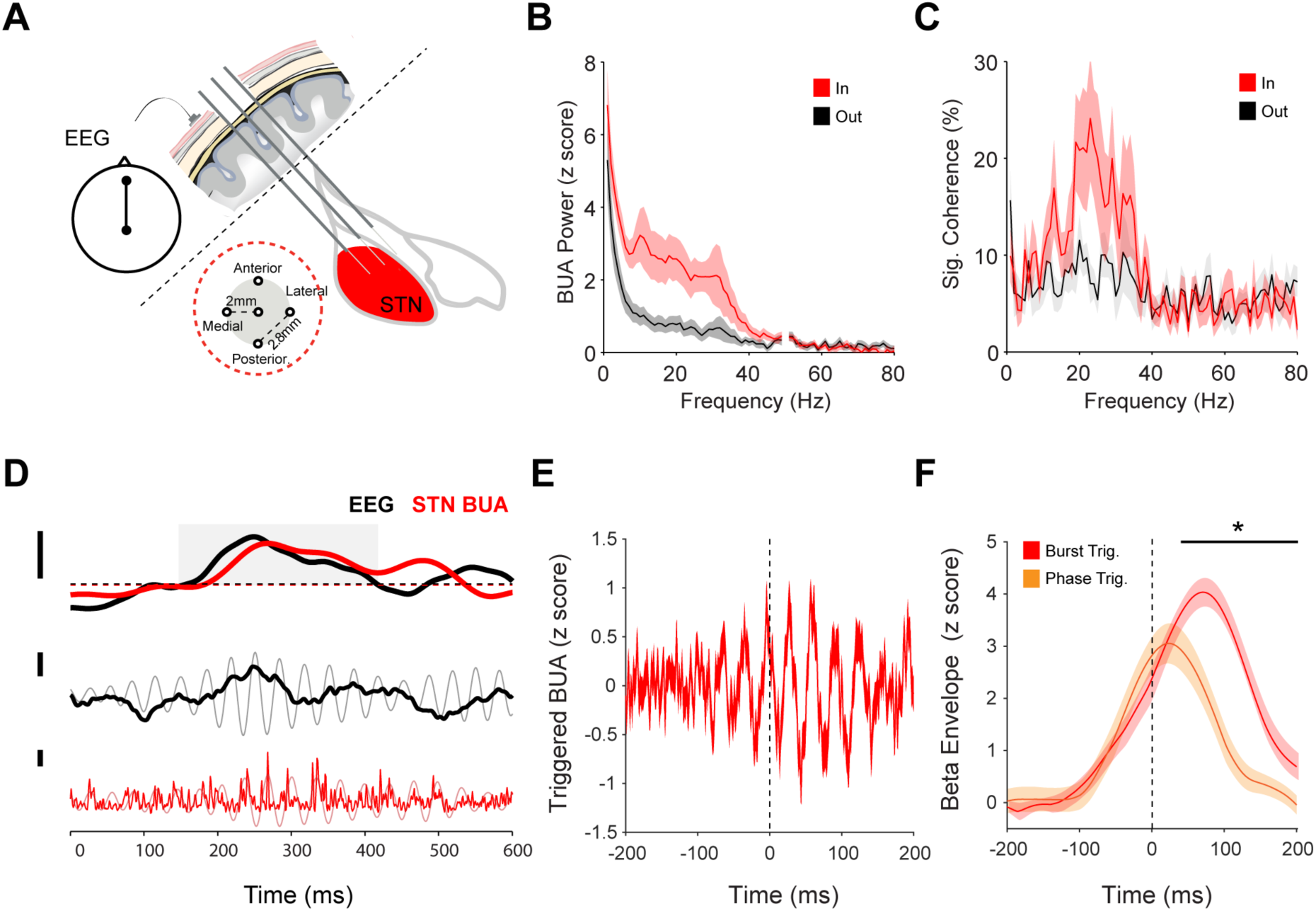
Characterizing cortically entrained beta bursts in the STN. **A**, Background unit activity signals were recorded from 5 microelectrodes in the STN area together with EEG from the Fz-Cz position. As the microelectrodes traversed the STN, some electrodes were inside and some outside the structure. **B**, Average BUA power spectrum from inside (red) and outside (grey) the STN, highlighting that enhanced rhythmic activity in the beta band is observed inside the STN. **C**, Over 80% of the BUA signals recorded inside the STN are coherent with the frontal EEG. **D**, An exemplary cortical beta burst from subject 1 (right hemisphere), together with simultaneously recorded STN-BUA. Dashed line indicates the amplitude threshold for determining the onset of the cortical beta burst. Onset of a cortical beta burst was defined as the time point cortical beta power exceeded the 75^th^ percentile of the cortical beta power and remained elevated for at least 50ms. The beta amplitude of both the EEG and STN BUA simultaneously crosses the burst threshold for around 200ms (top). This increase in amplitude is visible in the raw (grey) and filtered (black) EEG signal. Bursts of activity in the STN BUA (raw; red, beta filtered; pink) are clearly aligned to the trough of the EEG beta oscillation during the burst. **E**, STN-BUA exhibited increases in phase consistency at the onset of a cortical beta burst (subject 1, right hemisphere). **F**, The phase-aligned oscillations shown in E were enveloped using the Hilbert transform. Solid red line indicates the average z-transformed evoked BUA envelope averaged per subject across electrode contacts inside the STN (hemispheres = 13 patients = 7). Solid orange line indicates the average z-transformed BUA envelope evoked by cortical phase irrespective of the cortical beta power. Whether these envelopes were significantly different was determined using a cluster-based permutation test. Shaded regions indicate the standard error of the mean across all recordings

### Experimental Parkinsonism

#### In vivo electrophysiological recording with multielectrode arrays

Simultaneous extracellular recordings of unit activity and LFPs were made from numerous sites in the dorsal Str, GPe and STN of urethane-anesthetized 6-OHDA-lesioned rats. Striatal recordings were made without any other electrodes in the brain (n=7 animals, n=89 recordings). In some cases, STN and GPe recordings were made simultaneously (n=9 animals, n=25 recordings) and in other GPe was recorded alone (n=9 animals, n=19 recordings). Details on 6-Hydroxydopamine lesions and silicon probe recordings can be found in the Supplementary materials.

### Data Analysis

Recordings were analysed using a custom written software in MATLAB.

#### Deriving Background unit activity

For silicon probe recordings in experimental Parkinsonism, background unit activity was derived from raw probe recordings by high-pass filtering the recordings at 300 Hz using a third order Butterworth filter, followed by identifying spiking activity by setting a threshold at *mean+4sd* of the recording. A 4 ms segment was removed around each epoch crossing this threshold and replaced with part of the recording that did not contain any spiking activity. Following removal of spiking activity, the data was rectified, and low pass filtered at 300 Hz using a third order Butterworth filter. For STN microelectrode recordings from PD patients, the signal was recorded using a band-pass filter (300-6000 Hz). The other processing steps were identical.

#### Beta burst

Beta burst onset was derived from the motor cortex in 6-OHDA lesioned rat model of PD and from Fz-Cz EEG from PD patients. An epoch was labelled as a “beta burst” if the instantaneous cortical beta power exceeded the 75^th^ percentile for at least one beta cycle (i.e. 50 ms). Cortical beta burst onset (i.e. crossing the 75^th^ percentile threshold) was adjusted to the nearest (within ±30 ms of the threshold crossing) 0 degrees of the cortical beta phase unless otherwise stated. Cortical beta phase was derived from the angle of the Hilbert Transform.

#### Evoked BUA

In PD patients, EEG and BUA were down-sampled to 1000 Hz. Fz-Cz EEG was filtered between ±5Hz of the most coherent beta frequency (derived by averaging EEGBUA magnitude squared coherence from all recordings acquired inside the STN per hemisphere and subject) using a second order Butterworth filter. We (1) averaged unfiltered subthalamic BUA from each probe and recording depth, aligned to the onset of a cortical beta burst, (2) band-pass filtered each evoked BUA between ±5Hz of the most coherent beta frequency using a second order Butterworth filter, and (3) computed the z-score of the filtered evoked BUA. In 6-OHDA lesioned rat model of PD, we (1) averaged unfiltered BUA from each probe, aligned to the onset of a cortical beta burst, (2) band-pass filtered each evoked BUA between 15 and 35 Hz using a second order Butterworth filter, (3) computed the z-score of the filtered evoked BUA. If the maximum evoked BUA from a probe was less than 1.96 (zscore) (±200 ms around the burst onset) or if the summed cortico-subcortical coherence between 15-35 Hz was less than 10% of the total coherence between the two sites, evoked BUA from that probe was excluded from further analysis. The onset of change in phase consistency was defined as the first point when the envelope fitted over individual locked BUAs had a positive derivative. In order to ensure that onsets reflected the general trend in phase consistency, these signals were smoothed by a moving average filter of length 101 ms prior to determining the onset time points.

### 6-OHDA lesioned rat model of Parkinson’s disease

#### Power changes

Motor cortex and BUAs were down-sampled to 1000 Hz and filtered between ±5Hz of the most coherent ECoG-BUA beta frequency using a second order band-pass filter. Instantaneous power in the beta frequency band was derived from the Hilbert transform of the filtered signal. A moving average filter of length 50 ms was applied to the instantaneous power, since any power changes in the beta frequency band would occur at a much slower rate than the average period of the rhythm of interest (i.e. 50 ms). Changes in BUA_β_ power were assessed with respect to the median BUA_β_ power −500ms to 0ms prior to onset of a cortical beta burst. Whether these changes were significant was determined using a cluster-based permutation test, which compared the temporal consistency of subcortical BUA_β_ power change during cortical beta bursts to the modulation level of subcortical BUA_β_ power that was not locked to the onset of a cortical burst.

#### Single unit phase consistency

Spiking activity was discretized and represented as 1 that lasted for 1ms at the peak of the spike, while periods of non-spiking activity was represented as 0. In order to investigate the phase organization in single units, the discretized spiking activity was convolved with a Gaussian filter (reciprocal of standard deviation 25 ms) and averaged across all cortical beta bursts, following realignment to the onset of each cortical beta burst. We then band-pass filtered each burst-triggered spiking activity between 15 and 35 Hz using a second order Butterworth filter and computed the z-score of the filtered burst-triggered spiking activity. If a unit was not significantly phase locked to the cortical beta (Rayleigh Test, alpha=0.05), or the maximum burst-triggered spiking activity from a probe was less than 1.96 (z-score) (within ± 200 ms of burst onset), burst-triggered spiking activity derived from that unit was excluded from further analysis.

#### Single unit rate

Spiking activity was discretized and represented as 1 that lasted for 1ms at the peak of the spike, while periods of non-spiking activity was represented as 0. In order to investigate the phase organization in single units, the discretized spiking activity was convolved with a Gaussian filter (reciprocal of standard deviation 25 ms) and averaged across all cortical beta bursts, following realignment to the onset of each cortical beta burst. In order to differentiate between single unit phase consistency and rate, cortical beta burst onset was not phase-adjusted.

#### Phase slip

Instantaneous phase was derived from the Hilbert Transform of either the filtered BUA or the motor cortex. Phase alignment was computed by subtracting the BUA instantaneous phase from that of the motor cortex. Phase slips were computed by differentiating the unwrapped phase alignment between two regions and determining when z-scored difference between adjacent phase alignment values was greater than or less than 1.96. These points, referred to as phase slip, were represented as 1, while the other points were represented as 0. In order to compute the time evolution of phase alignment around a phase slip, the angular difference between the average phase alignment −50 to 0 ms prior to a phase slip and the average phase alignment 0 to 50 ms after the cortical beta burst onset was calculated. Similarly, the angular difference between the average phase alignment 0 to 50 ms after a phase slip and the average phase alignment 0 to 50 ms after the cortical beta burst onset was calculated.

## Results

The definition of a beta burst varies extensively across different studies, in terms of both, the threshold used to categorize an epoch as a beta burst, and the temporal dynamics of the identified burst (Feingold et al., 2015; Sherman et al., 2016; Tinkhauser et al., 2017a, 2017b). In this study, the occurrence of a cortical “beta burst” was inclusively defined as any period when EEG (PD patients) or frontal ECoG (experimental Parkinsonism) instantaneous beta amplitude exceeded the 75^th^ percentile of the cortical beta power across the recording (Tinkhauser et al., 2017b, 2017a). Bursts with a duration of 50 ms or less were excluded from further analysis due to ambiguity associated with the nature of transients that last less than one beta cycle (Cagnan et al., 2015). Subcortical spiking and background unit activity (BUA) were analyzed with respect to the defined onset and offset of cortical beta bursts, in order to provide a reproducible reference point for delineating neural dynamics within and across different BG structures. Nevertheless, the onset and offset a of beta burst defined as above are empirical definitions ‐as highlighted later, different processes precede this onset defined according to the 75^th^ percentile threshold crossing.

### Parkinson’s disease patients

#### STN - background unit activity temporal consistency changes at the onset of the cortical beta burst

Beta bursts have been extensively reported in the STN LFPs of PD patients, but it is unclear to what extent they reflect subthalamic unit activity. To address this issue, we utilized the background unit activity (BUA) signal, which provides an opportunity to investigate changes in local synchrony, and to explore the dynamics of a local ensemble of STN neurons. In line with subthalamic LFPs and single units in PD, BUA recorded from the STN of PD patients exhibited enhanced rhythmic activity in the beta frequency band (Fig. 1A, B). Moreover, subthalamic BUA was strongly coherent with the cortical activity in the beta frequency band (Fig. 1C). Beta oscillations and cortical synchronization were confined to BUA signals within the physiologically defined boundary of the STN (Fig. 1B, C).

In order to characterize the down-stream impact of cortical beta bursts, we explored their impact on the rhythmicity of subthalamic BUA. We observed that unfiltered STN-BUA exhibited increased phase consistency with respect to the cortical beta bursts (i.e. when instantaneous cortical beta power exceeded the 75^th^ percentile of the cortical beta power) (Fig. 1E). These results highlight that STN synchrony in the beta band has, on average, a fixed phase relationship with respect to the motor cortex, linking in oscillatory output of STN neurons. Critically, the locking of oscillatory STN-BUA began before the point that cortical beta power exceeded the 75th beta power threshold (Fig. 1E and 1F). The change in STN phase consistency began on average −131±99 ms (mean ± STD) prior to the cortical beta power exceeding the beta power threshold (hemispheres = 13 patients = 7 n = 18 recordings out of 232 recordings – please refer to Materials and Methods for further details). These findings demonstrate that, in PD patients, EEG and STN beta bursts reflect transient periods of oscillatory synchronization that will be propagated to other parts of the network by spiking activity.

### Experimental Parkinsonism

#### Subcortical BUA exhibits enhanced rhythmic activity in the beta band during a cortical beta burst

Electrophysiological recordings in the BG of PD patients are mostly limited to the STN. In order to define how other areas of the network are modulated by cortical beta bursts, we utilized recordings from the 6-OHDA hemi-lesioned rat model of PD, which allows for analyses of both ensembles of neurons and single unit activity of defined populations of neurons across the BG network (Mallet et al., 2008; Sharott et al., 2017). To establish that beta bursts in this model were similar to those in PD patients, we first analyzed the BUAs from the Str and GPe, as well as the STN. The evolution of BUA_β_ power across the BG was compared for coincident ECoG beta bursts that were separated into three durations (Fig. 2A). STN, GPe and Str BUA_β_ power increased with cortical burst duration; the longer the cortical beta burst lasted, the stronger the BUA_β_ became in all structures (Fig 2B). Whether these changes were significant was determined using a cluster-based permutation test, which compared the temporal consistency of subcortical BUA_β_ power change during cortical beta bursts to the modulation level of subcortical BUA_β_ power that was not locked to the onset of a cortical burst (i.e. with respect to baseline variability of subcortical BUA_β_ power). It should be noted that the average (−200 to 350 ms around the onset of cortical beta burst) surrogate BUA_β_ power changes were not different across the three subcortical nuclei (unpaired t-test between STN-GPe, STN-Str, GPe-Str with p = 0.0911, 0.0856, 0.6678). We also explored whether the BUA_β_ power became stronger with increasing cortical beta burst duration, as summarized per nucleus in Table 1. Similarly, the likelihood that a cortical and a subcortical beta burst coincided, depended on the duration of the cortical burst (Supplementary Figure 1). BUA_β_ power derived from a given probe contact inside the STN, GPe, and Str, exceeded the 75^th^ percentile of the corresponding BUA_β_ power 20-50% of cortical beta bursts as highlighted in Supplementary Figure 1. Cluster based statistics were applied −200 to 350 ms around the onset of cortical beta burst in order to ensure that epoch length difference did not confound the statistics. Unless otherwise stated, cluster significance was determined according to a of 0.05.

**Figure 2.**
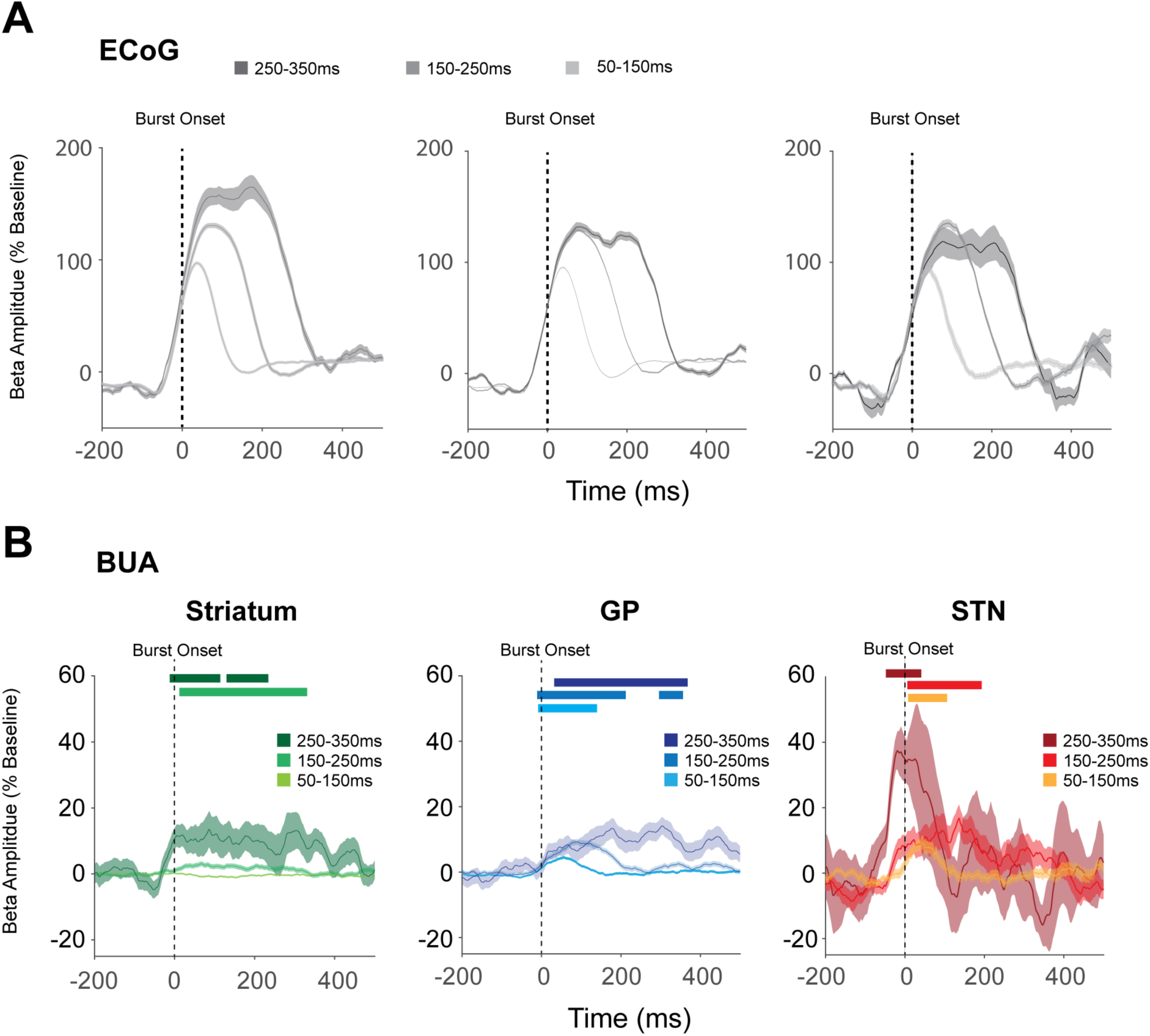
Str, GPe and STN BUA exhibit enhanced rhythmic activity in the beta frequency band during cortical beta bursts. **A**, The normalized amplitude envelope of the beta-filtered ECoG was aligned to the cortical beta burst onset, defined according to the 75^th^ percentile crossing. The onset of a cortical beta burst is indicated with a dashed line at time 0 ms. The resulting beta bursts were grouped into three durations. The three panels show the cortical beta bursts for the corresponding basal ganglia structures with which they were simultaneously recorded (aligned to the burst onset, defined according to the 75^th^ percentile crossing). **B**, The amplitude envelope of the beta-filtered BUA activity from the basal ganglia aligned to the cortical beta burst onset, defined according to the 75^th^ percentile crossing. Significant increases with respect to baseline are indicated with horizontal bars, color matched to the burst duration. Increases in BUA could reflect time locking or recruitment of a greater number of units to the burst onset. (Shaded regions indicate the standard error of the mean across all recordings made from a given structure)

**Table 1.** Timing of significant differences between BUA_β_ power changes observed during cortical bursts of different durations. Cluster significance corrected for multiple comparisons (α =0.0167 following Bonferroni correction).

|  | **50-150 vs. 150-250 ms** | **50-150 vs. 250-350 ms** | **150-250 vs. 250-350 ms** |
| --- | --- | --- | --- |
| **Str (n=848)** | 12-303 ms | -8-109 ms | no sig. cluster |
| **GPe (n=482)** | 25-209 ms | 111-351 ms | 174-351 ms |
| **STN (n=27)** | 121-180 ms | -46-26 ms | no sig. cluster |

#### Subcortical BUA is phase locked to the onset of cortical beta bursts

The above changes in BUA_β_ power suggest the overall recruitment of subcortical units is greater or units fire more consistently with respect to the cortical beta rhythm. In order to explore the link between subcortical BUAs and their phase relationship with respect to cortical beta, unfiltered BUAs were summed around the phase-adjusted burst onset, and then averaged across electrodes.

As the BUA signals in this analysis were unfiltered, modulation with respect to beta phase had to result from alignment of overall activity over trials. STN, GPe and Str BUAs exhibited increases in phase consistency with respect to cortical beta prior to the onset of a cortical beta burst (Fig. 3A). Increases in BG phase consistency preceded the time point when cortical beta power exceeded the 75^th^ percentile of cortical beta power. The increase in phase consistency began on average −159 ms (n=366), −117 ms (n=22) and −141 ms (n=199) in GPe, STN and the Str, respectively (Fig. 3B). The onset of change in phase consistency was defined as the first point when the envelope fitted over individual locked BUAs had a positive derivative. These results highlight that processes such as phase reorganization precede beta burst onset, as defined according to a threshold crossing.

**Figure 3:**
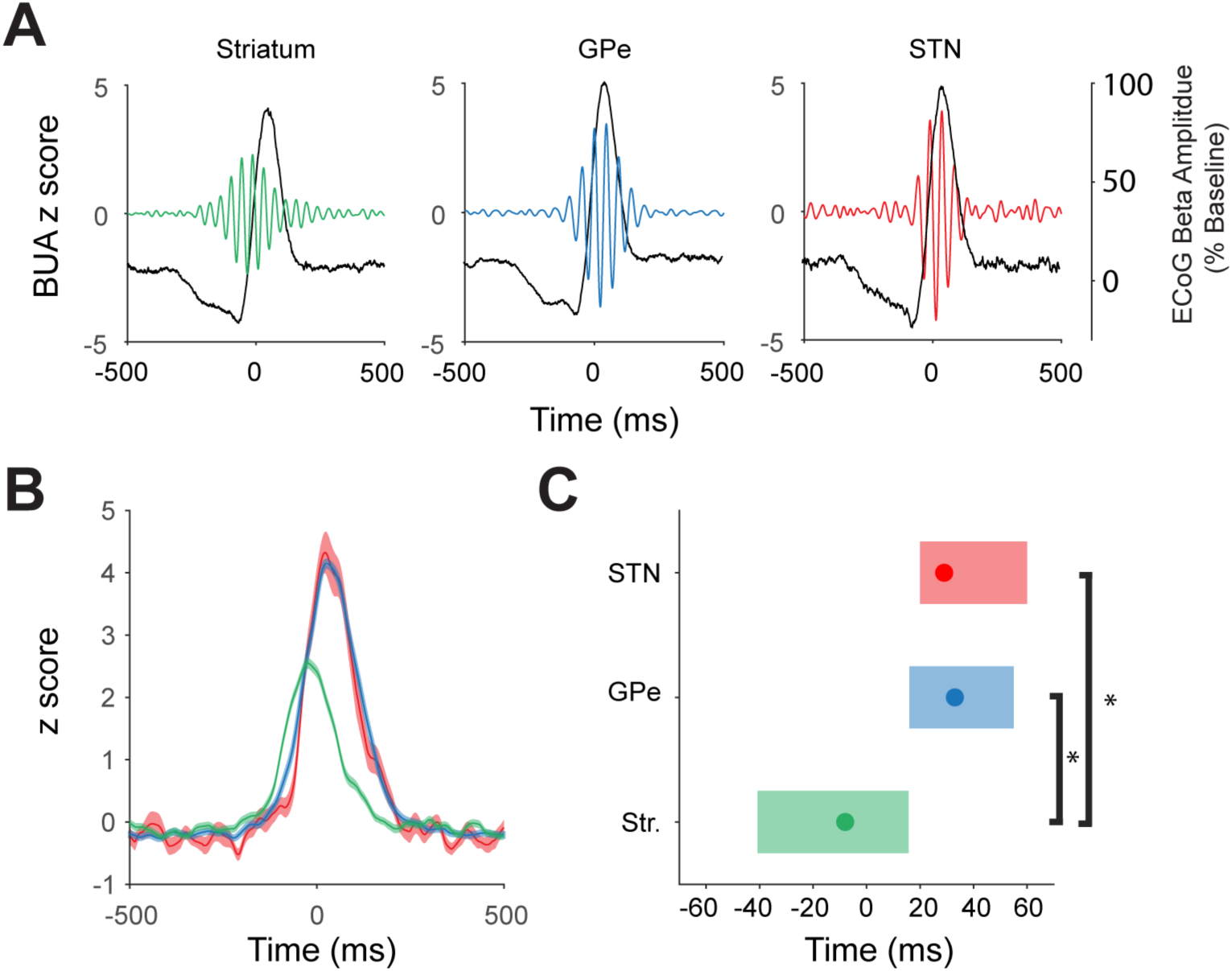
Subcortical beta emerges at a fixed phase relationship with respect to motor cortex. **A**, The STN, GPe and Str unfiltered BUA was averaged around the nearest cortical beta peak to the burst onset, such that zero-time always had the same cortical beta phase (colored lines, left y-axis). The same calculation was performed for cortical amplitude envelope (grey lines, right y-axis). In each structure, peaks and troughs in the amplitude of the unfiltered BUA signal, with a period of around 50ms, arose before and after the burst threshold indicating alignment with the cortical oscillation. **B**, The phase-aligned oscillations calculated in A were enveloped using the Hilbert transform. Solid lines indicate the average z-transformed evoked BUA envelope, while shaded regions indicate the standard error of the mean across all recordings made from a given structure. **C**, The median peak location per structure together with the 25^th^ and 75^th^ percentiles of the peak location: Str (n=199) and GPe (n=366) (Mann Whitney U test: p<0.001), and Str and STN (n=22) (Mann Whitney U test: p<0.001) peak locked BUA timings were significantly different. There was not a significant difference between the peak timing of the locked BUA in the STN and GPe (Mann Whitney U test: p=0.74).

The transient recruitment of Str to the on-going beta rhythm terminated prior to that of GPe and STN. The peak location for the evoked beta was significantly different between Str (n=199) and GPe (n=366) (Mann Whitney U test: p<0.001), and Str and STN (n=22) (Mann Whitney U test: p<0.001). There was not a significant difference between the peak locations of the evoked beta in the STN and GPe (Mann Whitney U test: p=0.74) (Fig. 3C).

#### Are relative increases in the amplitude envelope and phase consistency around the onset of beta bursts related to changes in firing rates of single neurons?

BUA provides an effective way of measuring recruitment of ensemble activity within a given structure by cortical activity. However, this signal could mask important differences in specific populations of neurons within those structures that have different baseline firing rates, patterns and phase relationships to cortical oscillations (Mallet et al., 2008, 2012). To address this, we performed the same analysis using sorted action potentials of single neurons and distinguished between the two major populations of neurons in the GPe, based on well-established criteria (Mallet et al., 2008). GP TI neurons fire at the inactive phase of the cortical slow oscillation and have been shown to correspond to prototypic GP neurons that project to STN. GP TA neurons fire at the active phase of the cortical slow oscillations and correspond to arkypallidal neurons that project only to Str (Mallet et al, 2012; Abdi et al, 2015). As GP neurons here were identified physiologically, based on the relationship to the cortical slow wave, we use the TI/TA nomenclature. Single unit firing rates in the three structures exhibited firing rates consistent with those previously reported. Specifically, Str neurons on average fired at 3 ± 4 spikes/sec (n=475), while STN (n=22), GP-TA (n=40), and GP-TI (n=179) neurons fired at 30 ± 13, 16 ± 10, and 12 ± 8 spikes/sec, respectively. Considering only the subset of neurons that were significantly locked to cortical beta, Str neurons on average fired at 5 ± 6 spikes/sec (n=104). As previously reported, the majority of these neurons were likely to be indirect pathway spiny projection neurons (Sharott et al., 2017). STN (n=18), GP-TA (n=33), and GP-TI (n=149) neurons fired at 32 ± 14, 16 ± 10, and 13 ± 7 spikes/sec, respectively.

The modulation of firing rate around cortical beta bursts was assessed by using burst-related time points as the trigger (time zero) around which to sum spikes over bursts, equivalent to peri-stimulus histogram. We first triggered unit activity by the amplitude-defined burst onset, without any alignment of that trigger to the same beta-phase over trials, which would detect slow changes in firing rate that aligned with the entire burst. This method did not result in any consistent modulation of firing rate in any of the BG populations (Fig. 4A), demonstrating that mean firing rate is not altered on the timescale of whole bursts. When the trigger was adjusted so that it occurred on the same phase for every burst (as in the BUA analysis), there were clear modulations in the firing of each population at beta frequency in line with the BUA analysis from each structure (Fig. 4B). Individual STN, GP TI, GP TA and Str units exhibited increases in their temporal consistency at the onset of cortical beta bursts (Fig. 4C). For individual GP and STN units, it is noteworthy that each beta burst (i.e. trial on the raster plot) could evoke a highly consistent pattern of oscillatory firing for over 200ms (Fig. 4C).

**Figure 4.**
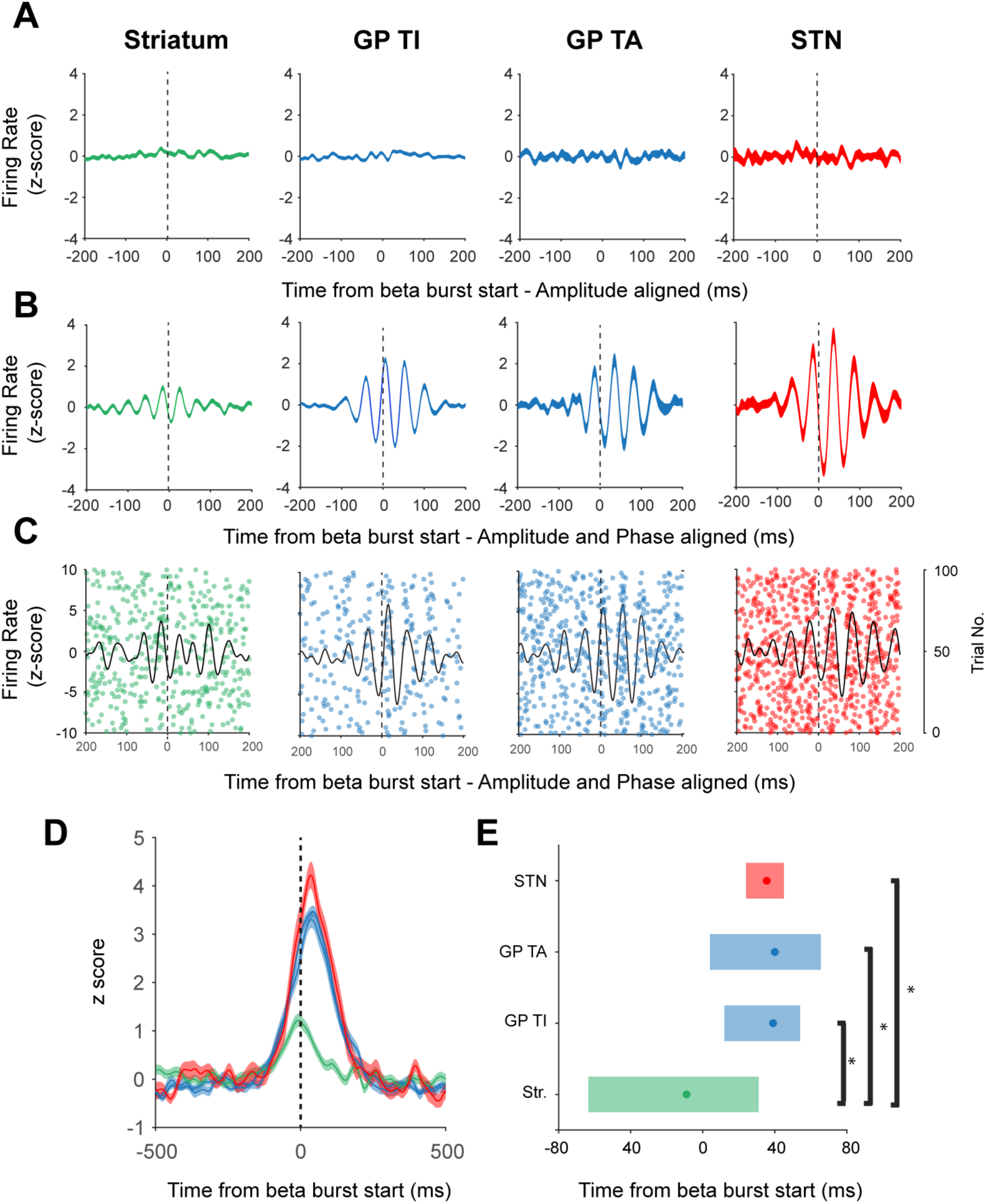
Basal ganglia units are recruited to the beta rhythm around the onset of a cortical beta burst. **A**, Mean z-score firing rate of the burst-triggered (time zero, 75^th^ percentile of amplitude) single units in each structure, with no adjustment for the cortical beta phase. Firing rate is not altered during the burst. **B**, The same analysis, but the burst-trigger is shifted to the nearest peak in the cortical beta phase. Firing rate is significantly modulated by the phase alignment to the cortical oscillation and offset with respect to time zero. **C**, Representative examples of unit activity from the Str, GP TA, GP TI and STN. Each sub-panel shows spike timings derived from a single unit, across individual cortical bursts. Black lines indicate the corresponding averaged activity for that unit realigned to the time point that cortical beta power crossed median levels. **D**, Lines indicate the average z-transformed, smoothed beta-envelope while shaded regions indicate the standard error of the mean across all recordings made from a given structure. **E**, The median peak timing per structure (dots) together with the 25^th^ and 75^th^ percentiles of the peak timing (shaded areas): The peak timing of the striatal units was significantly different from that of prototypic (Mann-Whitney U test: p < 0.001) and GP TA neurons (Mann-Whitney U test: p < 0.001) and subthalamic units (Mann-Whitney U test: p = 0.0034). In all plots, the onset of a cortical beta burst is indicated with a dashed line at time 0 ms.

In line with the BUA results, the local re-arranging of the precise timing of single unit discharges terminated first in the Str, and then in GP and STN (Fig. 4D). The peak location of the phase consistency of Str units (n=104) was significantly earlier than prototypic (n=149) (Mann-Whitney U test: p < 0.001) and arkypallidal neurons (n=33) (Mann-Whitney U test: p < 0.001) and subthalamic units (n=18) (Fig. 4E, Mann-Whitney U test: p = 0.0034). These results confirm that population dynamics derived from BUA are driven by changes in temporal consistency of spiking activity with respect to cortical beta phase and that this process begins prior to the amplitude-based burst threshold.

It is important to note that the differences in latency of cortical phase locking to not simply result from the different mean phase angles of spikes from a given neuron in relation to a single beta cycle, which have been reported previously (Mallet et al., 2008). Recruitment of striatal neurons was separated by that in STN and GPe by an average of 39 ms, when considering the BUA, and 47ms, when considering the spiking activity. If recruitment time was merely a confound of preferred spiking phase, then striatal, subthalamic and GP TA neurons should have exhibited comparable latencies in relation to the cortical beta burst, as they fire at the same part of the cycle. Indeed, such mean phase analysis cannot be used to infer temporal delay, as it is biased toward the center of the burst, where stable dynamics have been established and is determined partly by frequency (Sharott et al., 2018).

#### Phase reorganization takes place prior to a beta burst

Finally, we examined how phase alignment between cortical and subcortical BUA was altered around the onset of a cortical beta burst. We observed that, rather than synchronising gradually, phase alignment between cortical and subcortical nuclei altered abruptly – a phenomenon referred to as a phase slip. Converting phase alignment to a binary process whereby phase slips were denoted as 1 and other points were represented as 0, allowed us to compute the probability of observing a phase slip at a given time point across bursts. The likelihood of a phase slip being observed was higher prior to a cortical beta burst than during the burst (Fig. 5A). Slips could occur at multiple time points before the burst (Fig. 5A), irrespective of the burst length. The average probability of a phase slip in the 200ms prior to beta burst onset was greater than the average probability of a phase slip occurring in the following 200ms (paired t test p < 0.001 for all regions). In contrast, the time of phase slips following the burst was dependent on burst length (Fig. 5A).

**Figure 5:**
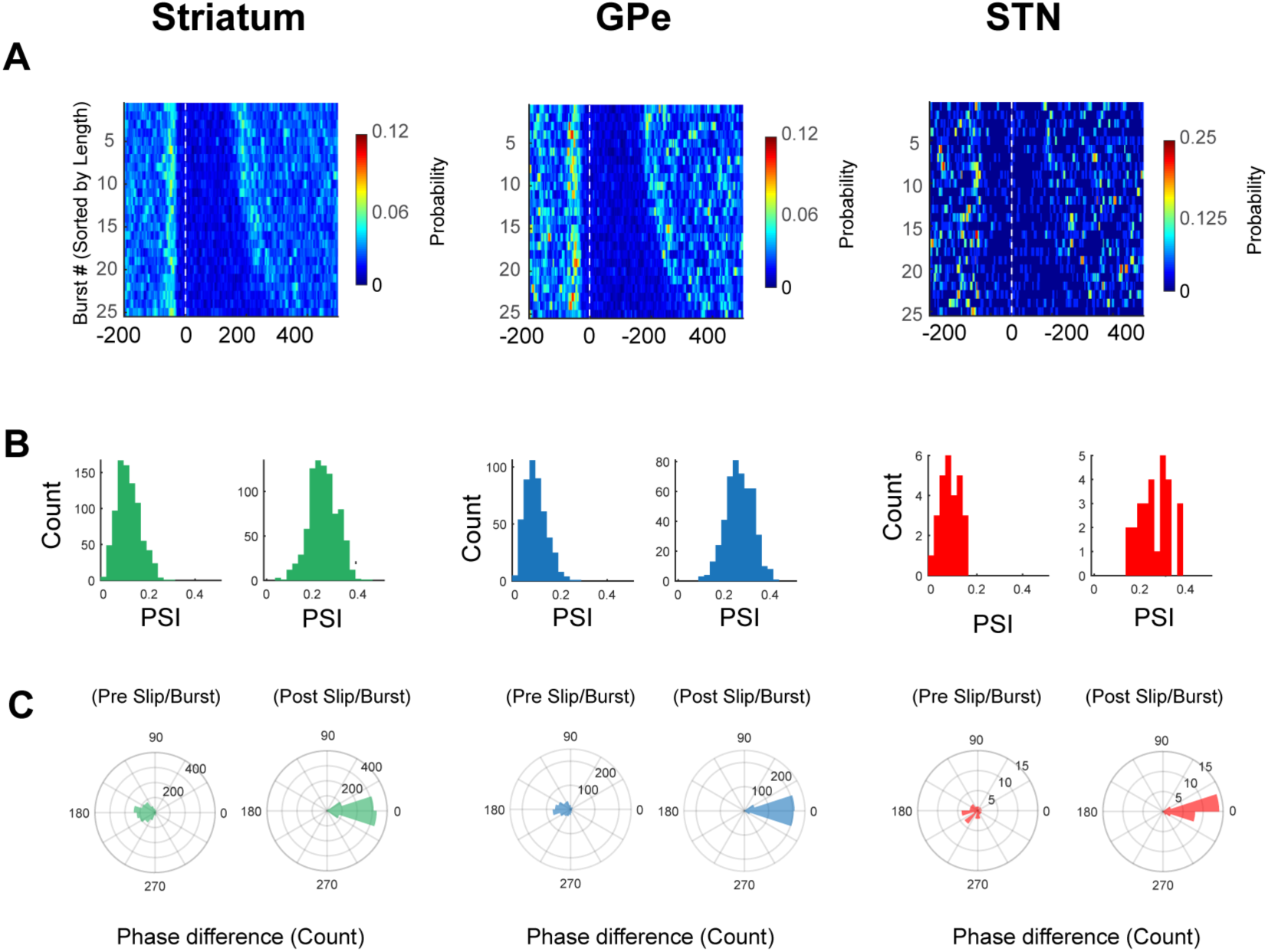
Phase alignment between the motor cortex and the basal ganglia nuclei changes prior to the onset of a cortical beta burst. **(A)** Average phase slips observed during the 25 longest cortical beta bursts in each recording. The phase alignment between the motor cortex and subcortical BUA altered prior to the cortical burst onset, defined according to the threshold crossing. **(B)** Phase alignment immediately prior to a phase slip differed from the phase alignment observed during a cortical beta burst. However, immediately after a phase slip, the phase alignment between two neural populations was the same as the phase alignment observed during a cortical beta burst (i.e. zero degrees difference between the phase of alignment after the slip and during the burst). **(C)** Phase alignment between the motor cortex and the basal ganglia nuclei, prior to a phase slip, was not consistent across different epochs-highlighted by low phase synchrony index (PSI) values. Post phase slip, phase alignment values were more consistent across different trials.

In order to determine the relationship of these phase slips to synchronisation of beta activities, we compared the properties of phase alignment before and after the slip occurred to that within the beta burst itself. We observed that the phase alignment between the ECoG and BUAs in all structures immediately before a phase slip was not consistent across different beta burst epochs, as revealed by low phase synchrony index (PSI) values (Fig. 5B). Immediately after a phase slip, the PSI values increased by more than double (Fig. 5B). Crucially, we found that the precise phase alignment occurring immediately after the phase slip was the same as that within the burst itself (Fig. 5C, as indicated by a difference of 0°). In contrast, there was a weak relationship between the pre-burst phase relationship and that within the burst, with a slight tendency for the opposite alignment (Fig. 5C, as indicated by a difference of 180°). These findings were repeated across all BG structures.

This analysis highlights that abrupt changes in the alignment of BG spiking activity to cortical beta oscillations can lead to phase relationships that result in the transiently stable oscillatory activity seen across the extended network in the form of beta bursts.

## Discussion

Our findings demonstrate the down-stream impact of cortical beta bursts on BG nuclei ‐Str, GPe and STN. These transient rhythmic activity patterns, which could last over 400 ms in PD patients and in 6-OHDA lesioned rats, were associated with subcortical synchrony and oscillation in single unit spiking in relation to the subsequent burst duration. Crucially, the interaction between the cortex and subcortical nuclei operated on two different time scales, one dictating the phase coupling across regions and the latter the extent of synchrony in the beta band, reflected as the spectral power of the beta oscillations. Phase reorganization of subcortical activity preceded the threshold crossing used for demarcating a cortical beta burst.

While previous studies on transient beta oscillations focused on the temporal dynamics of these events in a single brain region (Feingold et al., 2015; Sherman et al., 2016; Tinkhauser et al., 2017b), here, we characterized the downstream impact of beta bursts both on population synchrony and unit activity across the BG circuit. We observed that phase consistency of both subcortical BUA and spiking was altered, prior to the threshold crossing used for defining a cortical beta burst. The magnitude of phase consistency not only depended on the temporal coupling between the motor cortex and the subcortical nucleus, but also the spatial consistency of this phase reorganization across the subcortical brain region. These observations are supported by the theoretical models and experimental observations of Tass et al, who suggested that disrupting the spatial extent of synchrony inside the STN through coordinated reset could result in alleviation of PD symptoms (Adamchic et al., 2014; Tass and Majtanik, 2006; Tass et al., 2012).

### The Str and STN/GPe network couple to cortical beta bursts on different timescales

Exploring the transient dynamics across the motor circuit enabled us to examine the contributions of different brain regions to the on-going beta rhythm in PD. We observed that striatal recruitment to the cortical beta rhythm terminated prior to that in the STN and GPe. While inferences drawn from relative timings should be circumspect, as they may be complicated by different signal-to-noise levels, the BUA signal should be broadly equivalent across sites. The striatal neurons that contributed most heavily to the striatal BUA were likely to be indirect pathway spiny projection neurons (iSPNs), which become hyperactive following dopamine depletion (Mallet et al., 2006; Sharott et al., 2017). Importantly, the magnitude of hyperactivity of iSPNs is positively correlated with the strength of their phase locking to cortical beta oscillations (Sharott et al., 2017), suggesting that modulation of excitability is necessary for iSPNs to propagate cortical rhythms. The general hyperactivity of striatal neurons in intraoperative recordings (Singh et al., 2016), suggests that this mechanism could also apply in PD patients. In contrast, as firing of STN and GP neurons is driven by intrinsic currents, it is changes to timing and synchronization of their spiking that has a bigger impact on their output (Wilson, 2013). Such timing and synchronization would be influenced by the highly synchronized discharge of iSPNs (Sharott et al., 2017), which have a lower baseline firing rate, but are an order of magnitude greater in number.

Our results suggest that iSPN activity could propagate cortical oscillatory activity to STN and GP, which is then sustained beyond the initial striatal engagement. In this interpretation, the STN/GP network would thus serve as an amplifier of the corticostriatal oscillation in time. Conversely, experiments in MPTP-lesioned primates suggest that beta is propagated through the cortico-subthalamic “hyperdirect” pathway (Deffains et al., 2016, 2018; Tachibana et al., 2011). In the rat, however, highly rhythmic corticosubthalamic oscillations fail to pattern GPe unless dopamine is depleted, and/or striatal neurons are also coupled to the oscillation (Magill et al., 2001; Zold et al., 2012). The brief timescale of striatal engagement identified here could also explain the relative weakness of striatal beta oscillations in comparison to the STN in primates (Deffains et al., 2016). As the beta in primates is lower frequency and the burst duration is considerably longer (Deffains et al., 2018) than in rats and humans, the kind of analysis employed here might be necessary to reveal striatal involvement. Further work will be needed to delineate whether there is a genuine species difference in the way that parkinsonian oscillations are propagated, or that differences in methodology lead to the discrepancy in the proposed mechanism through which parkinsonian oscillations are generated and propagated.

### Phase reorganization and communication

The temporal relationship between individual units and neural populations influences not only the degree of information exchange, but also underpins transient changes in coupling between brain regions (Fries, 2005; Gerstner et al., 1996a, 1996b; Markram et al., 1997). For rhythmic neural activity, communication efficacy can be summarized through communication through coherence, which posits that there exists a certain angle between two populations that promote effective communication (Fries, 2015). It has been shown that the phase relationship between visual cortices are altered prior to an increase in coherence in the gamma band, confirming the mechanistic role of phase reorganization in facilitating neural communication (Womelsdorf et al., 2007).

Phase reorganization is facilitated by instantaneous changes in rhythmicity (i.e. frequency) of neural populations (Fries, 2015; Pelt et al., 2012). Context dependent modulations in cortical beta frequency has been observed over the motor cortex during cue expectancy and visuomotor preparation (Kilavik et al., 2012, 2013). In line with the communication through coherence theory, we observed that the phase alignment between the motor cortex and subcortical nuclei was transiently altered – a process referred to as a *phase slip*, prior to the onset of a cortical beta burst. Phase slips not only brought the phase alignment between two regions closer to the value associated with an increase in beta power, but also were observed at the end of a cortical burst i.e. when the cortical beta power decreased. These results highlight a neural process that could potentially facilitate modulation of beta power across the motor circuit and provide a complementary biomarker that could be used to control the timing and amplitude of stimulation delivered using DBS (Cagnan et al., 2015). In the context of PD, stimulation delivery could be timed according to the phase alignment observed across the motor circuit – for instance between the subthalamic and cortical neural activities. Critically, such a stimulation strategy could readily be implemented utilising the existing neuromodulation technologies (Swann et al., 2018), although it would require simultaneous recordings from several sites.

### Beta bursts and Parkinson’s disease

Beta oscillations have been extensively studied in the context of PD. Changes in beta power due to dopaminergic medication or during DBS have been associated with symptom severity (Eusebio et al., 2011, 2012; Kühn et al., 2006, 2009). Timing delivery of high frequency stimulation to STN according to the instantaneous beta power achieved greater symptom suppression than continuous high frequency stimulation (Little et al., 2013, 2015). Tinkhauser and colleagues (2017) addressed the paradox that delivering less stimulation can lead to greater efficacy by suggesting that such adaptive DBS terminated pathologically long bursts, while continuous DBS suppressed all beta oscillations, regardless of the burst duration (Tinkhauser et al., 2017b). In a follow-up study, the same group observed that therapeutic effects of dopaminergic medication are associated with termination of long duration subthalamic bursts and that patients’ symptom severity was positively correlated with the probability of long duration bursts (Tinkhauser et al., 2017a). Here, we characterized the transient changes in neural dynamics across the motor circuit. Changes in the magnitude of rhythmic activity patterns in the beta band in subcortical nuclei scaled with cortical burst duration ‐the longer the cortical beta burst lasted, the stronger the beta band activity became in subcortical structures. It should be noted that the probability of observing coincident beta bursts across the motor circuit and the BG also increased with cortical burst duration. These results further highlight the importance of beta burst duration while determining when and how much to stimulate deep brain structures in PD (Little et al., 2013, 2015). Critically, when we assessed the changes in subcortical firing rate around a cortical beta burst, we did not observe a significant change. These results highlight that the increase in the strength of the beta oscillations is primarily driven by the changes in the timing of the neural activity and in this instance, changes in the timing are not driven by changes in the rate of the neural activity. It has been previously observed that dopaminergic medication terminates phase coupling between the STN and globus pallidus that promotes further exaggeration of beta oscillations (Cagnan et al., 2015). By highlighting the effect of transient synchronisation of spiking activity across the BG network, the findings presented here provide further evidence that instantaneous temporal alignment across structures can profoundly influence activity widespread populations of neurons for 100s of milliseconds. Given that the duration of synchronisation appears to be the most relevant metric for manifestation of motor symptoms in PD (Deffains et al., 2018; Tinkhauser et al., 2017b), driving stimulation based on temporal alignment of network has the potential allow the disruption of pathological synchronisation at an earlier time point than the oscillation amplitude. Such an approach could lead to increased specificity and effectiveness of closed-loop DBS strategies.

## Acknowledgements

The authors would like to thank all the patients who participated in this study.

## Supplementary Materials

### Intraoperative recordings

#### Patient details and clinical scores

Please refer to Sharott et al for preoperative patient selection criteria (Sharott *et al.*, 2018). Patient details and clinical scores, together with medication dosages, are detailed in Supplementary Table 1.

#### Surgical procedures for PD patients

Please refer to Sharott et al for surgical details (Sharott *et al.*, 2018). The surgical target was determined using a combination of an MRI-compatible frame (Stryker Leibinger, Freiburg, Germany) and individual computerized tomography scan with T1 and T2 weighted MRI. A commercially available algorithm was used to fuse different neuroimaging modalities (iPlan, BrainLAB Inc., Westchester, IL, USA). Fused images were used to define the STN/nigra complex, the anterior/posterior commissure, and blood vessels. The surgical target (i.e. STN) was defined as 11-13 mm lateral to the midline, 1-3 mm inferior and 1-3 mm posterior to the mid-commissural point on both hemispheres. Electrode trajectories used for microelectrode recordings aimed to avoid sulci, blood vessels and ventricles.

#### Experimental Parkinsonism

Adult male Sprague Dawley rats (Charles River) were used for all experimental recordings. All experimental work adhered to Society for Neuroscience Policies on the Use of Animals in Neuroscience Research and were in agreement with the Animals (Scientific Procedures) Act, 1986 (United Kingdom). Animals were housed in a temperature-controlled environment with a 24hr light/dark cycle and *ad libitum* access to food and water.

#### 6-Hydroxydopamine lesions of midbrain dopamine neurons

Please refer to the following papers for details on 6-hydroxydopamine (6-OHDA) lesions (Mallet *et al.*, 2006, 2008a, b, 2012; Abdi *et al.*, 2015). All surgical procedures were performed during the day in a dedicated surgery room. 6-OHDA lesions took place under anesthesia. Anesthesia levels were assessed regularly by testing reflexes to a cutaneous pinch. In order to lesion midbrain dopamine neurons, 1 μl of 6-OHDA solution was injected 4.1 mm posterior and 1.2–1.4 mm lateral of Bregma, and 7.9 mm ventral to the dura (Paxinos *et al.*, 2007). Following surgery, animals were closely monitored and assessed for recovery using several metrics. Only animals that were judged to have recovered sufficiently were assessed for the severity of lesion and used for electrophysiological recordings. Lesion efficacy was assessed 15 days after 6-OHDA injection. The lesion was classified as effective when animals performed ≥80 contraversive rotations in 20 min following apomorphine administration (0.05 mg/kg, s.c.; Sigma) (Abdi *et al.*, 2015).

#### Silicon Probe recordings

Electrophysiological recordings were performed 21–39 days after 6Hydroxydopamine lesions of midbrain dopamine neurons under anesthesia. Details on silicon probe recordings have been previously described (Magill *et al.*, 2006; Mallet *et al.*, 2008a). High impedance unity-gain operational amplifiers were used to record monopolar signals from each probe contact (Advanced LinCMOS; Texas Instruments). A screw above the contralateral cerebellum was used as reference. Signals (ECoG and probe) were each sampled at 16.7 kHz using a Power1401 amplifier and Spike2 software from Cambridge Electronic Design Limited. Anesthesia levels were assessed by examination of the ECoG and by testing reflexes to a cutaneous pinch. Following electrophysiological recordings, animals were euthanized. A fixative was used for transcardial perfusion (Magill *et al.*, 2006; Mallet *et al.*, 2008a).

The probe signals were bandpass filtered between 500 and 6000 Hz. Single-unit activity was extracted according to the following criteria: 1) signal/noise ratio of >2.5 and 2) spike sorting using methods such as template matching, principal component analysis, and supervised clustering (Spike2 Cambridge Electronic Design Limited). Units were classified as single units if a distinct refractory period in the inter-spike interval histograms could be identified. Spiking activity recorded from the GPe was assigned to Type A (TA) or Type I (TI) populations based on whether they preferentially discharged on the positive “active” or negative “inactive” component of the cortical 1Hz oscillation during the slow wave brain state (Mallet *et al.*, 2008a) using activation histograms (Mallet *et al.*, 2008a), visual inspection of the raw data and the lag of the central peak in the spike triggered average of the cortical ECoG. Spike trains that did not fit unambiguously into the TA or TI group were not considered further in this study.

---

**Fig. S1:**
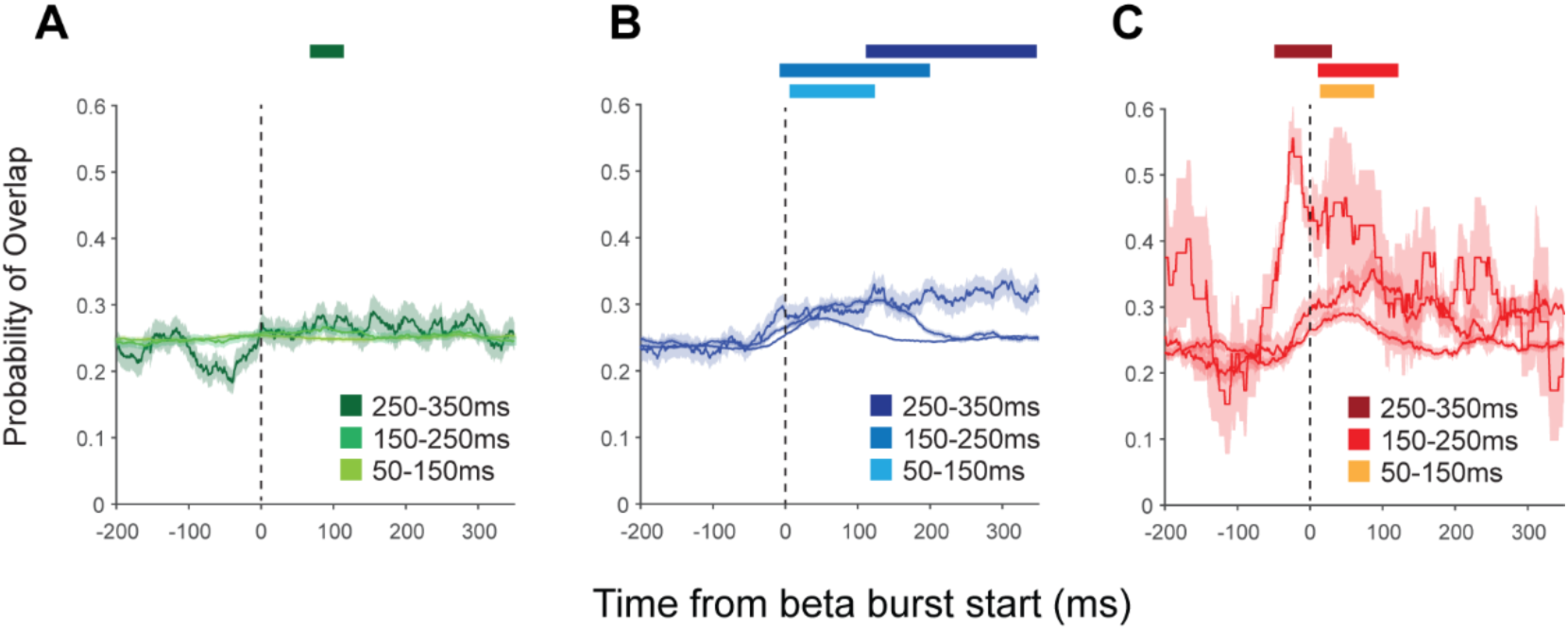
The likelihood for a cortical beta burst and a subcortical beta burst to coincide. The probability that a cortical beta burst coincided in time with a beta burst in the Str, GPe and STN increased with cortical burst duration. In order to assess the changes in burst overlap probability, subcortical instantaneous beta power was represented as one when it exceed the 75th beta power threshold, determined individually for each subcortical contact, and as 0 when it did not exceed the threshold. These were subsequently aligned to and averaged across different cortical beta bursts in order to obtain the likelihoods for cortical beta bursts to coincide with the subcortical ones. (Shaded regions indicate the standard error of the mean across all recordings made from a given structure)

**Fig. S2:**
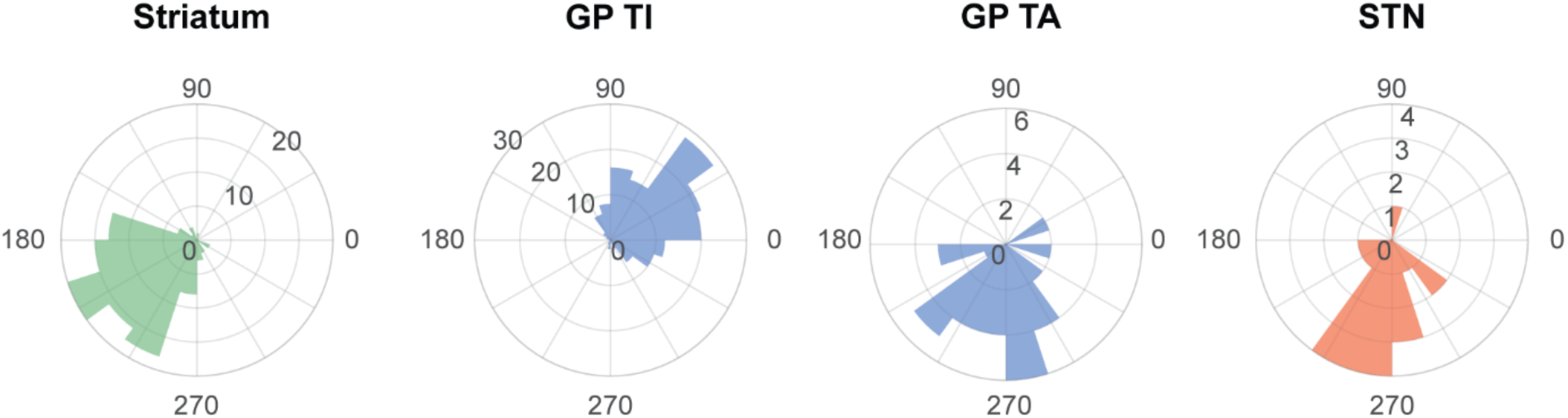
Mean phase locking angles of basal ganglia neurons in cortical beta oscillations. Histograms of the mean angles of phase locking for single units from the different basal ganglia neuronal populations.

**Supplementary Table 1.** Patient Details and Clinical Scores.

| Case | Age (Yrs) and Sex | Disease Duration (Yrs) | Motor UPDRS OFF | Motor UPDRS ON | Medication pre-operation | Hoehn/Yahr Score | Dominant side | Major Symptoms | Hemispheres analysed |
| --- | --- | --- | --- | --- | --- | --- | --- | --- | --- |
| 1 | 61F | 25 | 50 | 30 | Levodopa 700 mg<br>Carbidopa 150 mg<br>Entacapone 1000 mg<br>Benserazide 25 mg<br>Pramipexol 0.26 mg | 5 | Left | Tremor,<br>Akinesia/rigidity<br>Fluctuations | Right |
| 2 | 70F | 8 | 38 | 21 | Ropinirole 8mg<br>Alpha-dihydroergocryptine 40 mg<br>Amantadine 600 mg<br>Levodopa 250mg<br>Carbidopa 150mg | 3 | Left | Bradykinesia<br>Fluctuations<br>Dyskinesia | Both |
| 3 | 67M | 25 | 55 | 19 | Levodopa 1250 mg<br>Entacapone 1400 mg<br>Carbidopa 312.5mg<br>Rotigotine 6 mg<br>Amatadine 150mg | 3 | Right | Equivalence<br>Fluctuations<br>Dyskinesia | Both |
| 4 | 64M | 15 | 53 | 39 | Amantadine 300 mg<br>Levodopa 550 mg<br>Ropinirole 20 mg<br>Entacapone 900 mg<br>Carbidopa 112.5 mg<br>Benserazide 25 mg | 4 | Left | Akinesia/rigidity<br>Camptocormia | Left |
| 5 | 69F | 9 | 21.5 | 7 | Levodopa 450 mg<br>Lisuride 0.9 mg<br>Rotigotine 4 mg<br>Amantadine 300 mg | 3 | Right | Equivalence type<br>Fluctuations<br>Dyskinesia | Both |
| 6 | 66M | 11 | 28 | 14 | Amantadine 150 mg<br>Levodopa 1450 mg<br>Tolcapone 300mg<br>Carbidopa 312.5<br>Benserazide 50mg | 4 | Right | Akinesia/rigidity<br>Fluctuations | Both |
| 7 | 70F | 26 | 45 | 24 | Entacapon 1600 mg<br>Amantadin 300 mg<br>Pramipexol 0.8755 mg<br>Madopar 125 T (Levodopa<br>100 mg + Benserazid 25 mg)<br>8x0.5<br>Nacom 100 retard<br>(Levodopa 100 mg +<br>Carbidopa 25 mg) 1x1<br>Selegilin 5 mg | 4 | Left | Tremor,<br>Akinesia/rigidity | Left |

